# A climate-resilient microalga, *Desmodesmus ecotolerans* sp. nov., for biophotovoltaic applications

**DOI:** 10.64898/2026.09.24.754046

**Authors:** Hyeonsik Yoon, Joong-Tak Yoon, Mengwei Cheng, Shaban Muhammad, Thomas Li, Zixu Han, Jaeyeon Lee, Jungbin Kim, Ho-Seok Lee, Jangyong Kim, Eun Yu Kim

**Author notes:** Corresponding authors. E-mail addresses (H.-S. Lee),;(J. Kim) (E.Y. Kim). These authors contributed equally to this work and share first authorship (Hyeonsik Yoon, Joong-Tak Yoon).

## Abstract

Microalgae are promising biological platforms for sustainable energy production because they can convert photosynthetic activity into electricity in biophotovoltaic (BPV) systems. Identifying strains that grow well at elevated temperatures and under reduced irradiance is an important step toward BPV applications in seasonally variable environments. Microalgae were isolated from Seondong Lake (Seoul, Republic of Korea) during three sampling periods and screened for growth at different temperatures. Among 45 isolates, KHUC-11 showed the highest weighted growth index based on growth rates at 25, 30, and 35 °C under dark conditions and was selected for further characterization. Phylogenetic reconstruction of the 45S rRNA gene cluster, internal transcribed spacer 2 (ITS2) secondary-structure comparison, and morphological characterization supported the description of KHUC-11 as *Desmodesmus ecotolerans* sp. nov. *D. ecotolerans* produced more biomass than *Chlamydomonas reinhardtii* CC-125 under both moderate- and low-light conditions. Under low light, its final dry biomass concentrations were 1.81 and 1.58 times those of CC-125 under mixotrophic and photoautotrophic conditions, respectively. Despite its smaller cell size, *D. ecotolerans* showed higher photosynthetic performance than CC-125, and screen-printed electrode (SPE)-based electrochemical assays demonstrated enhanced anodic current generation and CV-derived estimated power output. Taken together, these findings identify *D. ecotolerans* as a novel climate-resilient *Desmodesmus* species with bioelectrochemical properties that support further evaluation for BPV applications.

## 1. Introduction

Renewable energy technologies that reduce carbon emissions while maintaining energy supply are urgently needed. Microalgae can convert solar energy and atmospheric CO_2_ into biomass and value-added products, while offering rapid growth, high photosynthetic efficiency, and cultivation on non-arable land. [1-8] Microalgae have been investigated for applications, including wastewater treatment, carbon capture, food and feed production. [9-13] Microalgae can also serve as living photosynthetic catalysts in biophotovoltaic (BPV) systems for electricity generation. [6, 14] Despite these advantages, practical deployment of microalgal technologies remains limited by cultivation costs and unstable performance under fluctuating environments. [14-16] Many strains evaluated under controlled laboratory conditions may not maintain comparable productivity in outdoor or semi-outdoor systems, where temperature and irradiance vary over daily and seasonal timescales. [17] This gap between laboratory performance and field applicability remains a major bottleneck for algal bioenergy technologies.

A promising strategy to address this limitation is to explore environmental isolates from local habitats. Unlike laboratory model strains maintained under stable conditions, environmental microalgae may have been naturally selected under fluctuating temperature, irradiance, nutrient, and microbial conditions. Such isolates may therefore represent a valuable yet relatively underexplored reservoir for discovering robust strains capable of maintaining productivity under realistic cultivation conditions. [17]

This concept is particularly relevant in East Asian monsoon climates, including Seoul, where algal cultivation can be exposed to seasonal temperature shifts, summer heat, winter cold, monsoon rainfall, frequent cloud cover, and variable solar irradiance. Such fluctuations can reduce growth stability, photosynthetic performance, and biomass productivity in outdoor or semi-outdoor cultivation systems. Therefore, microalgal strains that maintain growth and physiological activity under temperature and irradiance fluctuations may provide advantages for bioenergy applications in seasonally variable regions.

BPV systems use photosynthetic microorganisms to convert light-driven metabolism into electrical output, making photosynthetic performance and extracellular electron transfer important traits when evaluating candidate strains. [14, 18] Because BPV devices may be exposed to variable irradiance and daily or seasonal temperature fluctuations, strains that sustain photosynthesis and electrochemical output under these conditions may improve long-term device performance.

To identify microalgal resources suitable for renewable-energy applications under seasonally variable conditions, environmental microalgae were collected from Seondong Lake, a small artificial freshwater lake on the Kyung Hee University Seoul campus, Republic of Korea, and subjected to seasonal sampling and temperature-based screening. This screening identified KHUC-11 as the highest-performing isolate across 25–35 °C. Phylogenetic analysis of the 45S rRNA gene cluster, internal transcribed spacer 2 (ITS2) secondary-structure comparison, and morphological characterization supported its description as a novel species, *Desmodesmus ecotolerans* sp. nov. We further compared *D. ecotolerans* KHUC-11 with the model green alga *Chlamydomonas reinhardtii* CC-125, a well-characterized reference strain previously used in a microphotosynthetic power cell [19]. The comparison included growth performance, biomass accumulation, photosynthetic activity, and screen-printed electrode (SPE)-based electrochemical output. These analyses were used to evaluate *D. ecotolerans* as a climate-resilient bioenergy microalga with potential for biophotovoltaic applications.

## 2. Materials and Methods

### 2.1 Culture media and growth conditions

Microalgal strains were maintained in Tris-acetate-phosphate (TAP) medium [20]. Tris-phosphate (TP) medium was prepared by omitting acetate from the same formulation, whereas nitrogen-depleted TAP (TAP-N) medium was prepared by omitting the nitrogen source while retaining all other components of TAP medium. The pH of all media was adjusted to 7.0 before use. For strain purification, TAP medium was supplemented with ampicillin (700 μg mL^−1^), cefotaxime (200 μg mL^−1^), and carbendazim (1 μg mL^−1^). [21] Growth assays were conducted in TAP or TP medium under continuous illumination at either 50 or 120 μmol photons m^−2^ s^−1^. Cultures were acclimated for 3 days under the corresponding experimental conditions and inoculated at an initial optical density at 750 nm (OD_750_) of 0.05. Growth assays, except for temperature-screening experiments, were conducted at 25 °C under continuous illumination.

### 2.2 Collection and isolation of microalgal strains

Environmental microalgae were collected from Seondong Lake, Kyung Hee University, Seoul, Republic of Korea (37.5948° N, 127.0501° E), in August, October, and November 2024. Samples were spread onto TAP agar plates and repeatedly purified on antibiotic-supplemented TAP agar until unialgal isolates were obtained. Isolates from August, October, and November were designated KHUA, KHUB, and KHUC, respectively. *Chlamydomonas reinhardtii* CC-125 was used as the reference strain.

### 2.3 Temperature-based growth screening and selection of candidate isolates

All isolates were cultivated in TAP medium at 25, 30, 35, and 40 °C without illumination. Growth was monitored for 14 days by measuring OD_750_. Maximum specific growth rates (μ) were calculated from the exponential phase of growth curves. To facilitate strain prioritization across temperature-screening conditions, a weighted growth index (WGI) was developed for this study and calculated as follows:

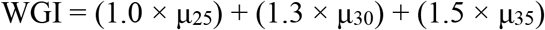

where μ_25_, μ_30_, and μ_35_ represent the maximum specific growth rates at 25, 30, and 35 °C, respectively. Isolates with WGI > 2.0 were selected for molecular characterization.

### 2.4 Amplification and sequencing of the 45S rRNA gene cluster

Genomic DNA was extracted using Chelex-100 resin. [22] The nuclear 45S rRNA gene cluster was amplified using the primer pair ss5/LR3 and sequenced by Sanger sequencing (Bionics, Republic of Korea). Primer information is provided in Table S1. Assembled sequences were queried against the NCBI GenBank database using BLASTn and subsequently used for phylogenetic analysis and secondary-structure analysis of internal transcribed spacer 2 (ITS2). The KHUC-11 sequence was deposited in GenBank under accession number PZ597142.

### 2.5 Phylogenetic and ITS2 secondary-structure analyses

Reference sequences retrieved from GenBank included BLASTn matches and sequences from representative taxa within the Scenedesmaceae. The 18S rRNA, ITS1, 5.8S rRNA, ITS2, and 28S rRNA regions within the KHUC-11 45S rRNA gene cluster were annotated using ITSx with default settings. [23] Multiple sequence alignments were generated using MUSCLE. [24] The best-fit nucleotide substitution model was selected in MEGA12 based on the Bayesian information criterion (BIC), and the K2+G+I model was used for maximum-likelihood analysis. Phylogenetic relationships were reconstructed using maximum likelihood (ML), minimum evolution (ME), and neighbor-joining (NJ) methods implemented in MEGA12. [25] Bootstrap support values were estimated from 1,000 replicates. [26] ITS2 secondary-structure analyses were performed using LocARNA with default settings. [27] Compensatory base changes (CBCs), hemi-compensatory base changes (hCBCs), and sequence divergence were determined from pairwise sequence–structure alignments between KHUC-11 and *Desmodesmus abundans*.

### 2.6 Morphological characterization and imaging

KHUC-11 and CC-125 were characterized using fluorescence microscopy and holotomographic imaging. Bright-field imaging was performed only for KHUC-11. KHUC-11 cells were harvested during the exponential growth phase from cultures grown in TAP medium and imaged using an LSM 800 confocal laser scanning microscope (Carl Zeiss, Germany). Chlorophyll autofluorescence was recorded using the AF680 channel.

For holotomographic analysis, KHUC-11 and CC-125 were cultivated in TAP medium for 3 days or in nitrogen-depleted TAP (TAP-N) medium for 9 days. Cells were imaged using an HT-X1 holotomography system (Tomocube, Republic of Korea). Neutral lipids were fluorescently labeled with BODIPY 493/503 (Thermo Fisher Scientific, Cat. No. D3922). A 10 μM BODIPY 493/503 working solution was prepared using dimethyl sulfoxide (DMSO), and harvested cell pellets were resuspended directly in the staining solution, resulting in a final BODIPY concentration of 10 μM. Samples were incubated for 50 min at room temperature. [28]

Following staining, samples were centrifuged at 700 × *g* for 5 min, the supernatant was discarded, and the cell pellet was resuspended in phosphate-buffered saline (PBS) to remove excess staining solution. The washing step was repeated once under the same centrifugation conditions. The washed cell suspension was then adjusted to an OD_750_ of 0.200 by 2.5-fold dilution with PBS. For cell immobilization, 50 μL of the diluted cell suspension was rapidly mixed with 50 μL of 1% TAP-agarose, yielding a final agarose concentration of 0.5%, and the immobilized cells were subjected to holotomographic and fluorescence imaging.

For fluorescence acquisition using the HT-X1 system, chlorophyll autofluorescence was detected using the Cy5 channel, whereas BODIPY 493/503 fluorescence was detected using the FITC channel. Holotomographic reconstruction and subsequent image analysis were performed using TomoAnalysis Viewer software (Tomocube). Reconstructed refractive-index tomograms were used to examine three-dimensional cellular morphology, and cell size was quantified using the same software.

### 2.7 Biomass determination

Biomass accumulation was determined gravimetrically as dry cell weight (DCW) following standard microalgal biomass determination procedures. [5, 7] A defined culture volume (10 mL) was harvested by centrifugation at 500 × g for 5 min, and the cell pellet was washed three times with 1 ml of 0.5 M ammonium formate. The washed cells were filtered through pre-weighed GF/C filters, dried at 70 °C for 72 h, and weighed to determine the dry biomass. Biomass concentration was calculated from the increase in filter weight and expressed as dry cell weight per culture volume.

### 2.8 Photosynthetic and electrochemical analysis

Photosynthetic CO_2_ assimilation was measured using an LI-6800 Portable Photosynthesis System equipped with a 6800-18 Aquatic Chamber (LI-COR Biosciences, Lincoln, NE, USA). KHUC-11 and CC-125 cultures were adjusted to the same optical density (OD_750_ = 0.5) prior to measurement. CO_2_-response and light-response curves were generated by varying CO_2_ concentration and irradiance, respectively. CO_2_ assimilation rates were normalized to total chlorophyll content. Total chlorophyll content was determined from methanol extracts by measuring absorbance at 652, 665, and 750 nm and calculating chlorophyll concentrations according to Caccamo et al. (2024). [29]

For electrochemical measurements, cells were washed with 0.1× phosphate-buffered saline (PBS) and adjusted to the same optical density before loading onto gold screen-printed electrodes (SPE; 4-mm working electrode diameter; geometric area = 0.1257 cm2). A 40 μL aliquot was loaded onto the working electrode. Cyclic voltammetry (CV) was performed using a VIONIC potentiostat/galvanostat (Metrohm Autolab, Utrecht, The Netherlands) over a potential range of −0.2 to +0.6 V versus Ag/AgCl at a scan rate of 100 mV s^−1^. Blank-corrected current was calculated by subtracting the 0.1× PBS blank current from the sample current. Anodic current density, oxidative CV area, CV-derived estimated power output, and CV-derived estimated power density were calculated from blank-corrected voltammograms and normalized to total chlorophyll content loaded onto the electrode [30, 31].

### 2.9. Statistical analysis

Data are presented as mean ± SD from at least three biological replicates unless otherwise stated. Statistical analyses were performed using GraphPad Prism 10 (GraphPad Software, Boston, MA, USA). Differences between two groups were assessed using an unpaired two-tailed Student’s t-test, whereas multiple-group comparisons were performed using unpaired t-test with Welch’s correction, as appropriate. Differences were considered statistically significant at p < 0.05.

## 3. Results

### 3.1 Isolation and temperature-based screening of microalgal isolates

To identify temperature-adaptable microalgal strains for subsequent characterization, environmental isolates recovered from Seondong Lake (Seoul, Republic of Korea) were subjected to temperature-based growth screening. A total of 45 purified microalgal isolates were obtained (Fig. 1A), representing samples collected during three independent sampling periods and designated KHUA, KHUB, and KHUC. Temperature-based screening identified KHUC-11 as the highest-performing isolate among the 45 environmental isolates examined (Fig. 1B). KHUC-11 showed maximum specific growth rates of 0.975, 0.896, and 1.137 d^−1^ at 25, 30, and 35 °C, respectively, resulting in the highest WGI value of 3.845 (Fig. 1C–E). Based on this performance, KHUC-11 was selected for subsequent taxonomic, morphological, physiological, and electrochemical characterization.

**Fig. 1.**
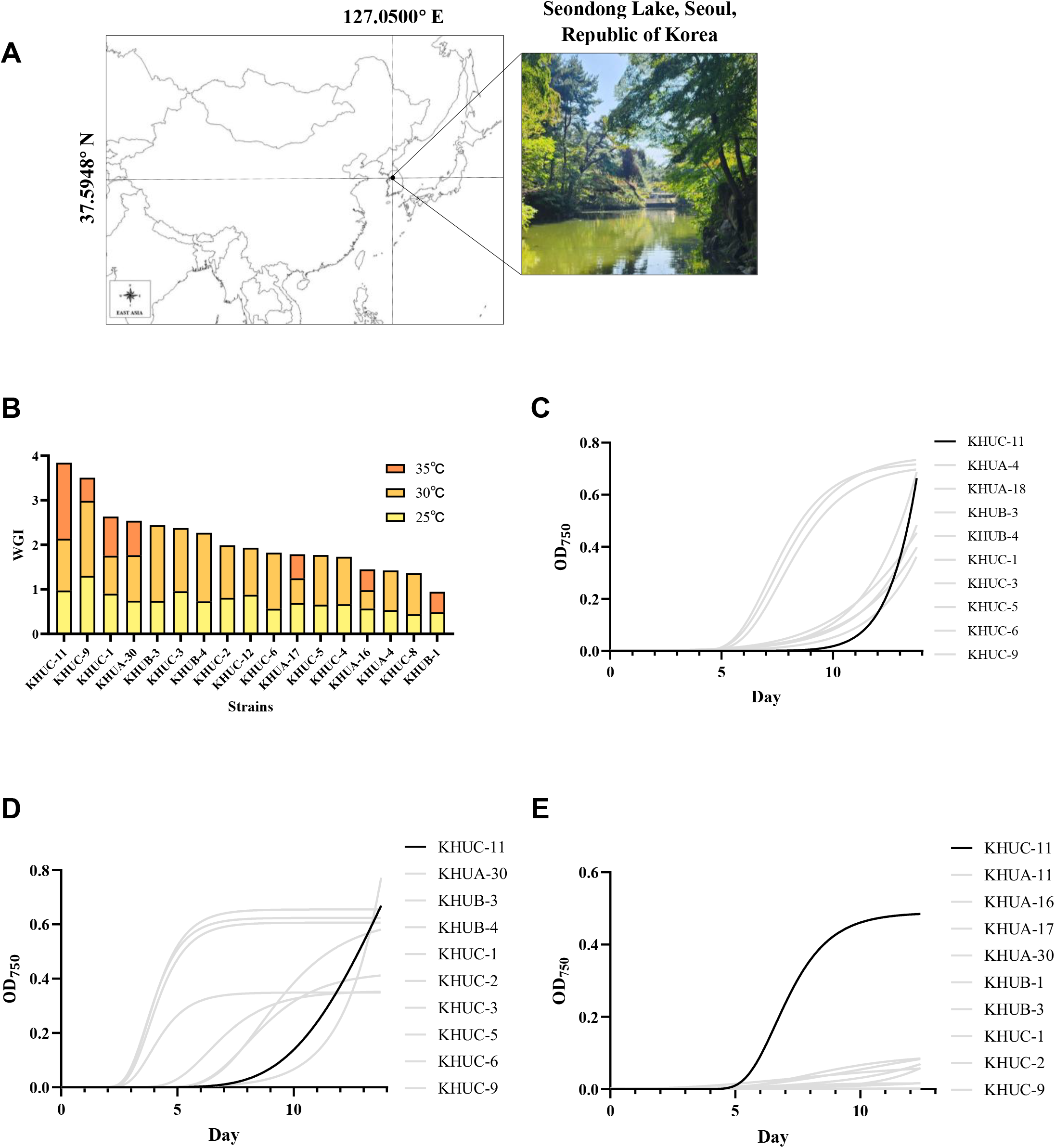
Collection and Screening of the samples from Seongdong lake. **Fig. 1A**. Geographical visualization of microalgae collection sites **Fig. 1B**. Result of thermotolerance screening **Fig. 1C**. Growth of KHUC-11 and other strains at 25°C **Fig. 1D**. Growth of KHUC-11 and other strains at 30°C **Fig. 1E**. Growth of KHUC-11 and other strains at 35°C

### 3.2 Molecular identification and taxonomic characterization of KHUC-11

To determine the taxonomic identity of KHUC-11, the nuclear 45S rRNA gene cluster was sequenced and subjected to phylogenetic and ITS2 sequence–structure analyses. Sequencing of the nuclear 45S rRNA gene cluster yielded a 3,428-bp sequence spanning the 18S rRNA gene, ITS1, 5.8S rRNA gene, ITS2, and partial 28S rRNA gene. BLASTn analysis identified *Desmodesmus opoliensis* as the closest database match, with 93.32% sequence identity and 85% query coverage. Phylogenetic analysis placed KHUC-11 within the genus *Desmodesmus* rather than the closely related *Scenedesmus* or *Tetradesmus* lineages (Fig. 2A). KHUC-11 was closely related to *Desmodesmus abundans* and formed an independent lineage supported by strong bootstrap values.

**Fig. 2.**
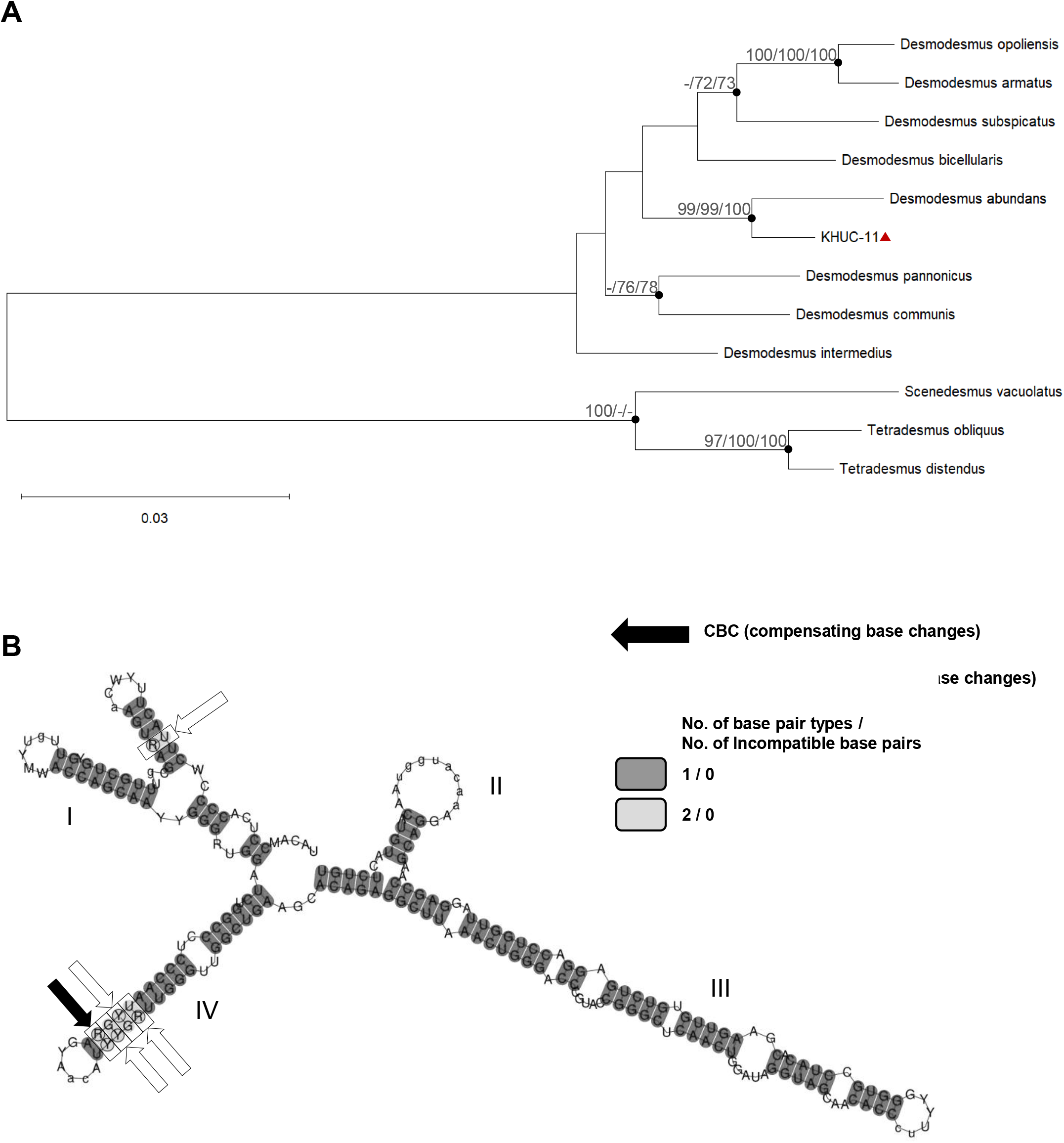
Phylogenetic analysis of KHUC-11 using 45S rRNA gene cluster DNA sequence. **Fig. 2A**. Phylogenetic tree constructed based on 18S rRNA, ITS1, 5.8S rRNA, ITS2, and 28S rRNA gene sequence. The selected DNA sequences were prepared by MUSCLE, and then operated MEGA 12 package programs such as Maximum Likelihood (ML), Minimum Evolution (ME), and Neighbor-Joining (NJ), while ML tree is representatively shown. 2318bp of the DNA sequence were used to construct the phylogenetic trees. The filled circle (•) indicates the branches which are simultaneously present in phylogenetic trees reconstructed by ML, ME, and NJ methods. Numbers at nodes indicates the bootstrap values of ML/ME/NJ based on 1000/1,000/1,000 replications. Bootstrap values higher than 70 are shown. KHUC-11 has triangle (▲) beside the taxon name. **Fig. 2B**. Secondary structures and sequence comparison of the internal transcribed spacer 2 (ITS2) sequence of KHUC-11 and *D. abundans*.

To further assess species-level differentiation from its closest phylogenetic relative, *D. abundans*, ITS2 secondary-structure analysis was performed. ITS2 secondary-structure comparison further distinguished KHUC-11 from *D. abundans* (Fig. 2B). The KHUC-11 ITS2 sequence was 236 bp, compared with 241 bp for *D. abundans*. One compensatory base change (CBC) and four hemi-compensatory base changes (hCBCs) were identified between the two taxa. Based on the phylogenetic and ITS2 sequence–structure analysis, KHUC-11 is proposed as a novel species, *Desmodesmus ecotolerans* sp. nov.

### 3.3 Morphological characterization of D. ecotolerans KHUC-11

To characterize the cellular morphology of *D. ecotolerans* KHUC-11, microscopic imaging was performed using bright-field, fluorescence, and electron microscopy. Microscopic imaging revealed that *D. ecotolerans* KHUC-11 cells were predominantly observed as single cells or paired cells and exhibited an oval to ellipsoidal morphology (Fig. 3A). In contrast, CC-125 cells were generally larger and displayed the characteristic morphology of unicellular *C. reinhardtii* cells. Chlorophyll autofluorescence was broadly distributed throughout the cell body, whereas DAPI staining revealed nuclear localization within individual cells. Cell-size analysis showed that KHUC-11 cells were significantly smaller than CC-125, with mean cell length of 5.386 ± 0.602 μm and width of 4.312 ± 0.495 μm, respectively (Fig. 3B). Holotomographic imaging further revealed the cellular architecture of KHUC-11, including the presence of thin cellular projections (Fig. 3C). These morphological characteristics were used for comparison with the reference strain CC-125 and subsequent physiological analyses.

**Fig. 3.**
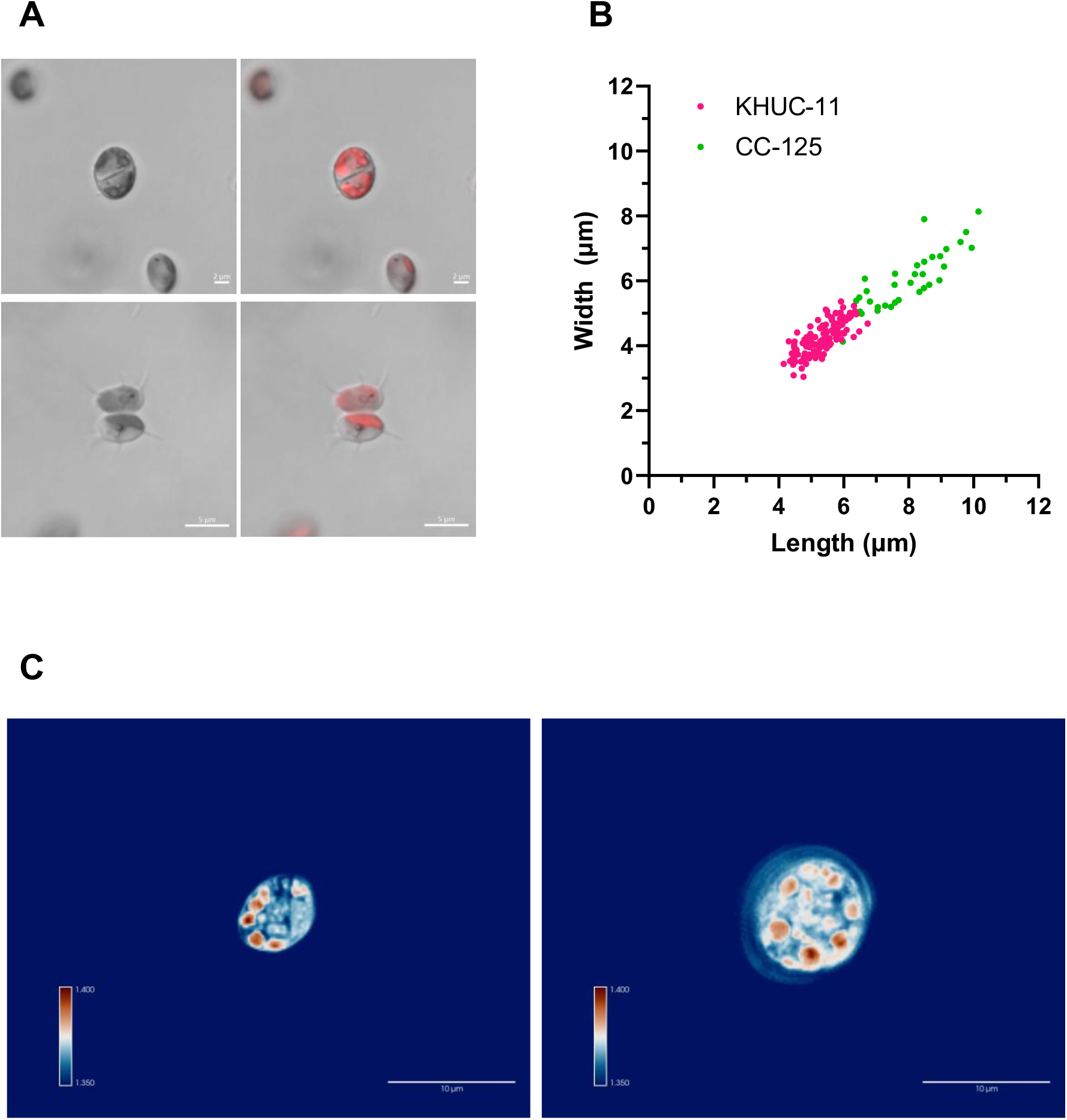
Morphological characterization of novel microalgae strains KHUC-11. **Fig. 3A**. Representative bright-field and chlorophyll auto-fluorescence KHUC-11 cells. Bright-field images show overall cell morphology. Scale bar = 5 um **Fig. 3B**. Quantitative comparison of cell size between CC125 and KHUC-11.. **Fig. 3C**. Holotomographic image of KHUC-11 (left) and CC-125 (right) showing detailed inner cell structure. Scale bar = 10 μm

### 3.4 Taxonomic assessment

Genus: *Desmodesmus* (R. Chodat) S.S. An, T. Friedl & E. Hegewald [32] Species: *Desmodesmus ecotolerans* H. Yoon, sp. nov.

Diagnosis: Cells occur predominantly singly or in pairs and are oval to ellipsoidal, measuring 5.386 ± 0.602 μm in length and 4.312 ± 0.495 μm in width. The chloroplast is cup-shaped, with a single pyrenoid-like region. *Desmodesmus ecotolerans* is distinguished from its closest phylogenetic relative, *D. abundans*, by its independent position in phylogenetic analyses of the nuclear 45S rRNA gene cluster and by one compensatory base change and four hemi-compensatory base changes in the ITS2 secondary structure.

Etymology. The specific epithet *ecotolerans* combines the modern combining form *eco-*, referring to environmental conditions, and the Latin present participle *tolerans*, meaning “tolerating” or “enduring.” The epithet was initially inspired by the thermotolerant phenotype of strain KHUC-11 identified during the temperature-screening assay.

Type locality: Seondong Lake, Kyung Hee University, Seoul, Republic of Korea (37.5948° N, 127.0501° E). Strain KHUC-11 was isolated from a sample collected on 5 November 2024.

Reference culture: Strain KHUC-11 is deposited in the Korean Collection for Type Cultures under accession number KCTC 16664BP and is maintained by cryopreservation in liquid nitrogen.

DNA sequence: Nuclear 45S rRNA gene cluster, 3,428 bp; GenBank accession number PZ597142.

### 3.4 Growth and biomass accumulation under different cultivation conditions

To compare growth performance under different trophic and irradiance conditions, KHUC-11 was compared with CC-125 under mixotrophic and photoautotrophic conditions at moderate- and low-light intensities. Under mixotrophic conditions, KHUC-11 reached higher final OD_750_ values than CC-125 under dim light conditions (Fig. 4A). Final OD_750_ values of KHUC-11 was low-light conditions, whereas CC-125 reached OD_750_ value of 1.94. Under photoautotrophic conditions, KHUC-11 also reached higher final OD_750_ values than CC-125 (Fig. 4C). Final OD_750_ value of KHUC-11 was 1.80 under low-light conditions, compared with 0.84 for CC-125. To determine whether the higher OD_750_ values of KHUC-11 were accompanied by greater biomass accumulation, dry biomass concentrations were measured.

**Fig. 4.**
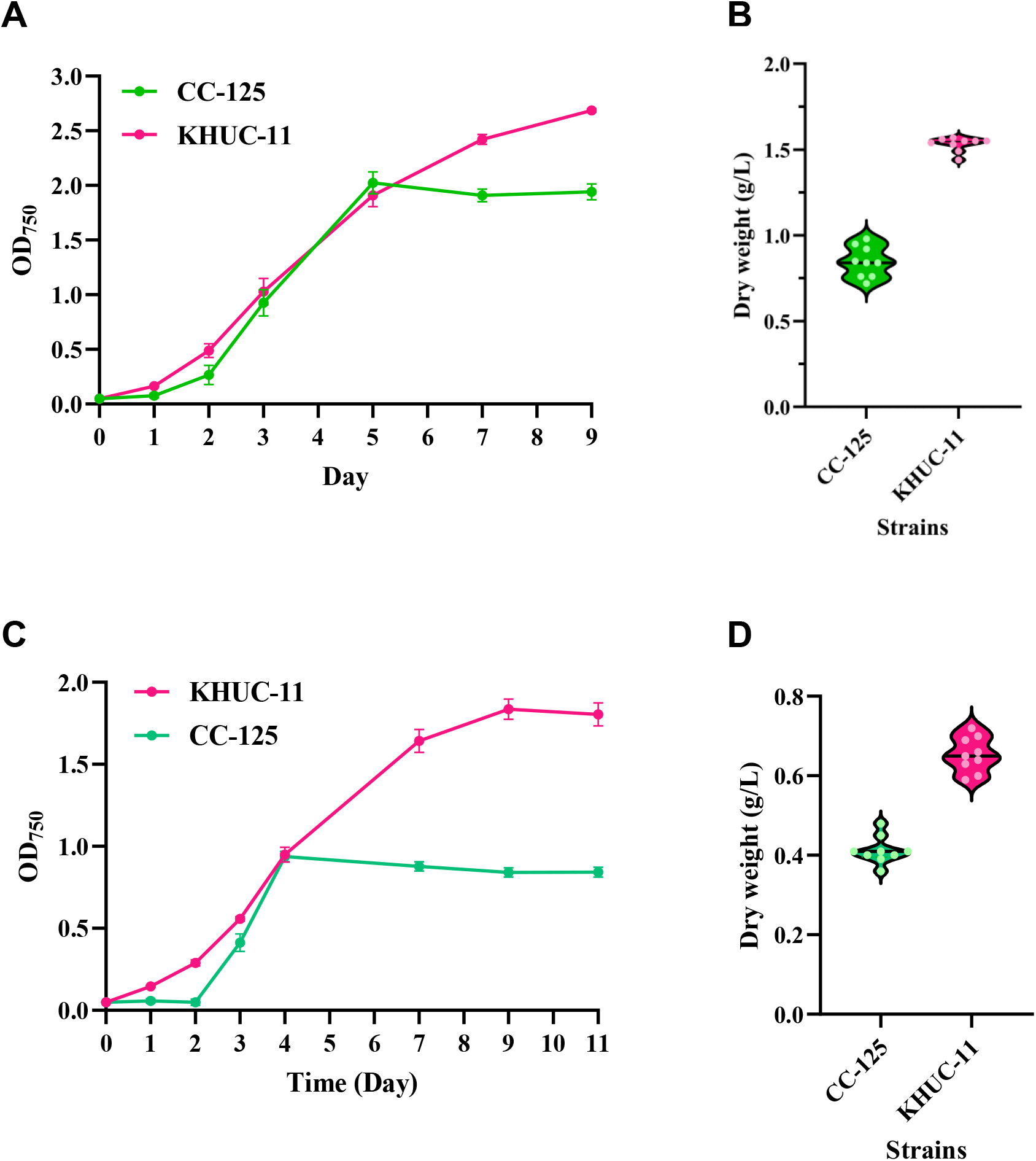
Growth comparison of CC-125 and KHUC-11. **Fig. 4A**. Mixotrophic growth curve under dim light condition **Fig. 4B**. Dry biomass accumulation of mixotrophic control under dim light condition **Fig. 4C**. Autotrophic growth curve under dim light condition **Fig. 4D**. Dry biomass accumulation of autotrophic control under dim light condition

Dry biomass measurements showed a trend consistent with the OD_750_ results. Under low-light mixotrophic conditions, KHUC-11 reached a final dry biomass concentration of 1.531 g L^−1^, compared with 0.847 g L^−1^ for CC-125 (Fig. 4B). Under low-light photoautotrophic conditions, KHUC-11 also accumulated more biomass, with a final dry biomass concentration of 0.653 g L^−1^, compared with 0.412 g L^−1^ for CC-125 (Fig. 4D).

### 3.6 Photosynthetic and electrochemical performance

To determine whether the enhanced growth of KHUC-11 was associated with photosynthetic and electrochemical activity, chlorophyll-normalized CO_2_ assimilation and SPE-based electrochemical analyses were performed. Chlorophyll-normalized CO_2_ assimilation analysis showed higher assimilation rates in KHUC-11 than in CC-125 across the tested CO_2_ concentration range (Fig. 5B). The maximum chlorophyll-normalized CO_2_ assimilation rate of KHUC-11 was 9.959 µmol µg^-1^ s^-1^ ×10^13^, compared with 4.166 µmol µg^-1^ s^-1^ ×10^13^ for CC-125. Light-response analysis also showed higher chlorophyll-normalized assimilation rates in KHUC-11 than in CC-125 across the tested irradiance range (Fig. 5C). The maximum chlorophyll-normalized assimilation rate under saturating irradiance was 13.750 µmol µg^-1^ s^-1^ ×10^13^ for KHUC-11 and 6.305 µmol µg^-1^ s^-1^ ×10^13^ for CC-125.

**Fig. 5.**
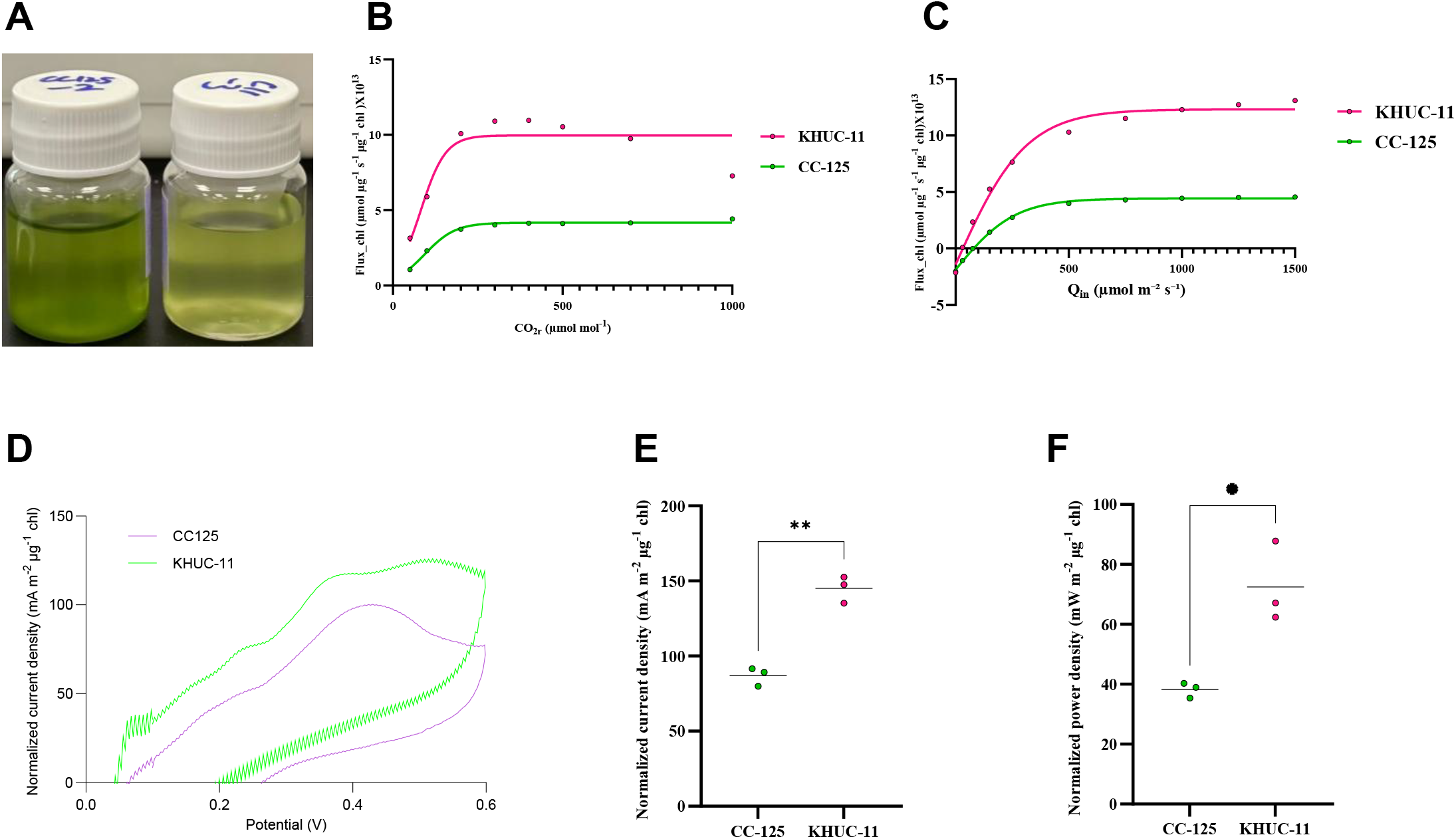
Comparative photosynthetic and electrochemical performance of CC-125 and KHUC-11. **Fig. 5A**. Representative images of CC-125 and KHUC-11. **Fig. 5B**. Carbon assimilation response curves measured under increasing CO_2_ concentrations. Carbon fixation rates were normalized to chlorophyll content and fitted using a non-linear saturation model. KHUC-11 exhibited a substantially higher carbon assimilation capacity than CC-125 across the tested CO_2_ range. **Fig. 5C**. Light-response curves showing photosynthetic carbon fixation rates under increasing incident light intensity. Data were normalized to chlorophyll content and fitted using a non-linear saturation model. KHUC-11 displayed a greater photosynthetic capacity and reached a higher light-saturated assimilation rate than CC-125. **Fig. 5D**. Cyclic voltammograms of CC-125 and KHUC-11 obtained using screen-printed electrodes. Current density values were normalized to chlorophyll content. KHUC-11 generated higher current densities throughout the scanned potential range, indicating enhanced extracellular electron transfer activity compared with CC-125. **Fig. 5E**. Maximum anodic current density derived from electrochemical measurements. Values were normalized to chlorophyll content and compared between strains. Each point represents an independent biological replicate, and horizontal lines indicate the mean. Statistical significance was determined using an unpaired two-tailed Welch’s t-test (** < 0.01). **Fig. 5F**. Maximum power density generated by CC-125 and KHUC-11, normalized to chlorophyll content. KHUC-11 exhibited significantly greater power output than CC-125. Each point represents an independent biological replicate, and horizontal lines indicate the mean. Statistical significance was determined using an unpaired two-tailed Welch’s t-test (* < 0.05).

To determine whether differences in photosynthetic performance were reflected in electrochemical output, SPE-based electrochemical analyses were subsequently performed. SPE-based cyclic voltammetry revealed higher chlorophyll-normalized current responses in KHUC-11 than in CC-125 throughout the scanned potential range (Fig. 5D). The maximum anodic current density of KHUC-11 was 145.10 mA m^-2^ μg^-1^ chl, compared with 86.92 mA m^-2^ μg^-1^ chl for CC-125 (Fig. 5E). CV-derived estimated power density was also higher in KHUC-11 than in CC-125 (Fig. 5F). The maximum estimated power density reached 72.43 mW m^-2^ μg^-1^ chl in KHUC-11, compared with 38.21 mW m^-2^ μg^-1^ chl in CC-125.

## 4. Discussion

### 4.1 Taxonomic significance of D. ecotolerans

The classification of KHUC-11 as a novel *Desmodesmus* species is supported by phylogenetic analysis, ITS2 sequence–structure comparison, and morphological characterization. Although phylogenetic analysis consistently placed KHUC-11 within the genus *Desmodesmus*, the isolate formed an independent lineage distinct from *D. abundans*, its closest phylogenetic relative (Fig. 2A). This taxonomic distinction was further supported by ITS2 sequence– structure analysis, which identified one CBC and four hCBCs between the two taxa. CBCs and hCBCs in ITS2 secondary structures have been widely used as informative markers for species delimitation in green microalgae and members of the Scenedesmaceae. ITS2 sequence– structure comparisons, including the identification of CBCs and hCBCs, have been widely used to distinguish closely related species within the Scenedesmaceae, including members of the genus *Desmodesmus*. [33] The phylogenetic and ITS2 sequence–structure analyses consistently distinguished KHUC-11 from its closest phylogenetic relative, *D. abundans*. Morphological features were consistent with characteristics of the genus *Desmodesmus*. KHUC-11 exhibited oval to ellipsoidal cells and characteristic surface projections observed by holotomography. Together, the phylogenetic, ITS2, and morphological evidence support the classification of KHUC-11 as a novel species, *Desmodesmus ecotolerans* sp. nov.

The specific epithet *ecotolerans* was initially inspired by the thermotolerant phenotype of KHUC-11 identified during the temperature-screening assay. Subsequent mixotrophic and photoautotrophic growth assays further showed that KHUC-11 maintained robust growth under reduced irradiance. Although elevated temperature and reduced irradiance were evaluated in separate experiments, these findings collectively provide broader ecophysiological context for the proposed epithet.

### 4.2 Growth and biomass production potential

A major objective of this study was to identify microalgal isolates capable of sustaining growth under cultivation-relevant stress conditions, including elevated temperature and reduced irradiance. Temperature-based screening identified KHUC-11 as the highest-performing isolate among the 45 environmental isolates examined. KHUC-11 consistently maintained robust growth across all tested cultivation conditions, which was reflected in increased biomass accumulation. In comparison with CC-125, KHUC-11 showed consistently higher growth performance under both mixotrophic and photoautotrophic conditions across moderate and low irradiance levels. These differences were further supported by dry cell weight measurements, confirming higher biomass accumulation in KHUC-11. KHUC-11 also maintained biomass accumulation under reduced irradiance, a condition relevant to natural outdoor environments influenced by cloud cover, seasonal variation, and self-shading in dense cultures. Overall, these results demonstrate that KHUC-11 exhibits sustained growth and biomass production across diverse environmental conditions, supporting its potential applicability for outdoor biomass production systems. Further validation under outdoor or semi-outdoor cultivation conditions will be required to confirm its scalability.

### 4.4 Photosynthetic and biophotovoltaic performance

To assess whether the enhanced growth performance of KHUC-11 was linked to improved photosynthetic and bioelectrochemical activity, photosynthetic performance and SPE-derived electrochemical responses were analyzed in parallel. Chlorophyll-normalized CO_2_ assimilation measurements revealed that KHUC-11 exhibited consistently higher carbon fixation rates than CC-125 across the tested CO_2_ concentration range, indicating improved photosynthetic efficiency on a pigment-normalized basis. Light-response analysis further confirmed enhanced photosynthetic performance of KHUC-11 under both moderate and low-light conditions, suggesting an improved capacity to maintain photosynthetic activity light-limited environments. In parallel, SPE-based cyclic voltammetry demonstrated that KHUC-11 generated higher chlorophyll-normalized current responses compared with CC-125 across the scanned potential range, indicating enhanced electrochemical activity at the cell–electrode interface. The increased anodic current density and larger CV-derived electrochemical output further support improved bioelectrochemical performance under the tested conditions. Collectively, the coordinated enhancement in photosynthetic CO_2_ assimilation and electrochemical response suggests functional coupling between metabolic activity and electron transfer processes in KHUC-11. This dual performance highlights its potential suitability for biophotovoltaic applications, particularly under low-light conditions relevant to natural and semi-outdoor environments.

## 5. Conclusions

This study identified a novel freshwater microalgal isolate, *Desmodesmus ecotolerans* KHUC-11, exhibiting robust growth across elevated temperature and low-light conditions relevant to outdoor cultivation environments. Compared with the reference strain CC-125, KHUC-11 demonstrated consistently enhanced growth performance and biomass accumulation under diverse cultivation regimes, supporting its physiological resilience under environmentally fluctuating conditions. Furthermore, integrated photosynthetic and electrochemical analyses revealed enhanced chlorophyll-normalized CO_2_ assimilation and increased current responses under SPE-based measurements, suggesting a functional coupling between metabolic activity and electron transfer processes. Overall, these results demonstrate that KHUC-11 represents a multifunctional microalgal strain with potential applications in biomass production and biophotovoltaic systems, under elevated temperature and low irradiance conditions relevant to outdoor environments.

## CRediT authorship contribution statement

Conceptualization: E.Y.K., J.K. (Jangyong Kim); Methodology: H.Y., J.-T.Y., M.C., S.M., T.L., Z.H., J.L., J.K. (Jangyong Kim), E.Y.K.; Investigation: H.Y., J.-T.Y., M.C., S.M., T.L., Z.H., J.L., J.K. (Jangyong Kim), E.Y.K.; Formal analysis: H.Y., J.-T.Y.; Data curation: H.Y., J.-T.Y., M.C., S.M., T.L., Z.H., J.L.; Visualization: H.Y.; Resources: J.K. (Jungbin Kim), H.- S.L., J.K. (Jangyong Kim), E.Y.K.; Supervision: H.-S.L., J.K. (Jangyong Kim), E.Y.K.; Project administration: E.Y.K.; Funding acquisition: H.-S.L., J.K. (Jangyong Kim), E.Y.K.; Writing – original draft: H.Y., E.Y.K.; Writing – review & editing: J.K. (Jungbin Kim), H.-S.L., J.K. (Jangyong Kim), E.Y.K.

## Declaration of competing interest

The authors declare that they have no known competing financial interests or personal relationships that could have appeared to influence the work reported in this paper.

## Funding

This work was supported by the Kunshan Science and Technology Bureau, China (grant no. kssc202302072 to E.Y.K.); the Research Fund for International Scientists (RFIS) of the National Natural Science Foundation of China (NSFC), China (grant no. 32350610246 to E.Y.K.); the Startup Fund of Duke Kunshan University, China (grant no. 00AKUG0122 to E.Y.K.); the Institute for Basic Science, Republic of Korea (grant no. IBS-R021-D1-2026-a00 to E.Y.K. and H.-S.L.); and the Research Development Fund of Xi’an Jiaotong-Liverpool University (grant no. RDF-24-02-067 to J.K.).

## Acknowledgements

The authors gratefully acknowledge the members of the CReAtE Lab for their valuable support and constructive discussions throughout this study.

## Data availability

The 45S rRNA gene cluster sequence of KHUC-11 has been deposited in GenBank under accession number PZ597142. Strain KHUC-11 has been deposited in the Korean Collection for Type Cultures under accession number KCTC 16664BP and is maintained by cryopreservation in liquid nitrogen.

## Declaration of generative AI and AI-assisted technologies in the writing process

During the preparation of this work, the authors used ChatGPT to assist with language editing. After using this tool, the authors reviewed and edited the content as needed and take full responsibility for the content of the publication.

## Notes

### Competing Interest Statement

The authors have declared no competing interest.

